# Single or Multi-Frequency generators in on-going brain activity: a mechanistic whole-brain model of empirical MEG data

**DOI:** 10.1101/084103

**Authors:** Gustavo Deco, Joana Cabral, Mark W. Woolrich, Angus B.A Stevner, Tim J. van Hartevelt, Morten L. Kringelbach

**Affiliations:** Theoretical and Computational Neuroscience Group, Universitat Pompeu Fabra, Barcelona, Spain; Institució Catalana de la Recerca i Estudis Avançats, Barcelona, Spain; Department of Psychiatry, University of Oxford, Oxford, UK; Center of Functionally Integrative Neuroscience, Aarhus University, DK; Oxford Centre for Human Brain Activity, University of Oxford, Oxford, UK; Centre for the Functional Magnetic Resonance Imaging of the Brain, University of Oxford, Oxford, UK

**Author notes:** Corresponding author: Joana Cabral, Department of Psychiatry, University of Oxford, Oxford, OX37JX, UK. These authors equally contributed for this work (Gustavo Deco, Joana Cabral). Conflict of interest: the authors declare to have no conflict of interest.

## Abstract

During rest, envelopes of band-limited on-going MEG signals co-vary across the brain in consistent patterns, which have been related to resting-state networks measured with fMRI. To investigate the genesis of such envelope correlations, we consider a whole-brain network model assuming two distinct fundamental scenarios: one where each brain area generates oscillations in a single frequency, and a novel one where each brain area can generate oscillations in multiple frequency bands. The models share, as a common generator of damped oscillations, the normal form of a supercritical Hopf bifurcation operating at the critical border between the steady state and the oscillatory regime. The envelopes of the simulated signals are compared with empirical MEG data using new methods to analyse the envelope dynamics in terms of their phase coherence and stability across the spectrum of carrier frequencies.

Considering the whole-brain model with a single frequency generator in each brain area, we obtain the best fit with the empirical MEG data when the fundamental frequency is tuned at 12Hz. However, when multiple frequency generators are placed at each local brain area, we obtain an improved fit of the spatio-temporal structure of on-going MEG data across all frequency bands. Our results indicate that the brain is likely to operate on multiple frequency channels during rest, introducing a novel dimension for future models of large-scale brain activity.

## 1 Introduction

Understanding the genesis of spatially and temporally structured brain rhythms is a crucial matter in neuroscience (Buzsáki, 2006, Wang, 2010, Womelsdorf et al., 2014). In vitro studies have shown that the cortical tissue is excitable, displaying the emergence of coherent oscillations under specific medium conditions while the single neurons fire only intermittently at lower (or higher) frequency (Buhl et al., 1998, Fisahn et al., 1998). Detailed computational models of spiking neurons have helped to investigate how neurons (and interneurons) connected in specific network topologies can generate firing patterns replicating electrophysiological measurements (Abbott and van Vreeswijk, 1993, Ermentrout et al., 2001, Brunel and Wang, 2003). The frequency of such oscillations is determined by time constants such as the feedback delay (relying on the excitatory-inhibitory loop), the synaptic time constants (delay and rise) and the axonal transmission times. Furthermore, the ratio of time scales of excitatory and inhibitory currents and the balance between excitation and inhibition also affect the properties of the rhythms.

To investigate how these locally generated oscillations interact at the macroscopic level of the whole brain network, it is useful to use neural-mass models in order to reduce the complexity of spiking neuron models to a small set of differential equations describing the population activity (Honey et al., 2007, Ghosh et al., 2008, Deco et al., 2009, Cabral et al., 2011, Cabral et al., 2014a). Following different reduction lines, these neural-mass models generate oscillations (with damped or constant amplitude) in a frequency tuned by the model parameters. Although these models have proved useful to investigate the source of long-range slow BOLD signal correlations, they assume a homogeneous oscillation frequency in every brain area, whereas evidence from EEG and MEG studies points in a different direction, i.e. that brain functional connectivity (FC) occurs in multiple frequency levels, displaying different functional networks according to the carrier oscillations (Mantini et al., 2007, de Pasquale et al., 2010, Brookes et al., 2011b, Hipp et al., 2012, Magri et al., 2012, Tagliazucchi et al., 2012, Keller et al., 2013, Hipp and Siegel, 2015).

The concept of a carrier oscillation implies that it is modulated (either through amplitude or frequency) by a ‘carried’ signal with much lower frequency. Indeed, several recent studies have detected meaningful long-range correlations in the slow amplitude envelopes (or power fluctuations) of narrowband neurophysiological signals obtained with EEG, MEG or LFP (Liu et al., 2010, Brookes et al., 2011b, Hipp et al., 2012, Magri et al., 2012, Engel et al., 2013, Keller et al., 2013, Hipp and Siegel, 2015, Brookes et al., 2016).

A number of models have aimed at replicating large-scale spontaneous electrophysiological data using large-scale network models with local neural-mass models, namely using the normal form of a Hopf bifurcation in the subcritical regime (Robinson et al., 2002, Freyer et al., 2011, Freyer et al., 2012) or simple Kuramoto oscillators (Cabral et al., 2014b). In these models, the cortico-thalamic and/or the cortico-cortical connectivity (with associated time-delays) serve as the structural support for a multistable regime, in which the system switches between the local node dynamics and global network dynamics with particular spectral properties.

In particular, Cabral and colleagues (2014b) have proposed that envelope correlations of alpha- and beta-band oscillations are originated by the metastable large-scale synchronization of locally generated gamma-band oscillations at 40Hz. This relies on the fact that, due to the presence of delays in the order of 10ms, the collective oscillations are more stable at reduced frequencies falling in the alpha and beta ranges (Niebur et al., 1991). Although this model provides an insightful mechanistic explanation for the phenomena observed in resting-state MEG, whether this mechanism actually occurs in the brain remains uncertain.

In this work we propose two novel candidate scenarios to explain the envelope correlations of carrier oscillations observed in the brain at rest. We address the problem from a different perspective where each brain area, as a complex neural mass operating in a critical regime, spontaneously generates oscillations on receiving sufficient input. Since the transition to the oscillatory state is induced by external noisy input, nodes receiving correlated input (through structural connections) are more likely to display the simultaneous emergence of oscillations, which in turn results in correlated envelopes shaped by the structural connectivity. Following empirical observations in resting-state MEG data where alpha- and beta-band oscillations emerge and dissipate in a correlated fashion throughout the cortex forming different functional networks, we remove the constraint that each neural mass can only resonate in the gamma-frequency band (30 to 100Hz) (in contrast with Cabral et al. (2014b)), and assume that brain areas may also resonate at slower frequencies (<30Hz) due both to complex internal connectivity and to external thalamo-cortical connectivity (Buzsáki, 2006, Palva and Palva, 2007, Engel et al., 2013). Moreover, we investigate two scenarios: one where local brain areas generate oscillations in a single fundamental frequency using the normal form of a Hopf Bifucation model; and another where each local brain area can resonate at multiple different frequencies (i.e. multiple Hopf bifurcation models, each with a different fundamental frequency). From a biophysical perspective, each brain area has millions of neurons coupled in complex connectivity patterns which may not unlikely resonate at a broad range of frequencies rather than a single one. In addition, cortical areas are connected to subcortical structures with intrinsic rhythmicity, such as the thalamus, which is believed to drive cortical oscillations in the alpha-frequency band (Lopes da Silva et al., 1997). As such, considering multiple rhythms within a brain area is not only biophisically plausible but should be considered in models of coupled brain areas to obtain a better picture of the whole-brain network dynamics captured with MEG.

The model outputs are compared with resting-state MEG brain activity, focusing on the temporal dynamics of envelope correlations as a function of the underlying carrier frequency. We follow the line of recent works from the fMRI and MEG literature that investigate the temporal dynamics of FC (Hutchison et al., 2013, Allen et al., 2014, Baker et al., 2014, Hansen et al., 2015), which have demonstrated that resting FC is definitively not static but shows a specific spatio-temporal structure, which is neglected when considering solely the static FC computed as an average over the whole recording time. Indeed, the so-called FCD (FC dynamics) allows for a better characterization of the data and a more accurate constraint on the model. We will show that the maximal richness of the MEG FCD is present around the carrier frequency of 10-16 Hz, in the same range where static band-limited amplitude correlations are known to be maximal in electrophysiological data (Brookes et al., 2011b, Hipp and Siegel, 2015, Siems et al., 2016).

Considering the whole-brain model with a single frequency generator in each brain area, we obtain the best fit with the empirical MEG data when the oscillation frequency is tuned at 12Hz. However, when multiple frequency generators are placed at each local brain area, we find the model largely outperforms the single-frequency case, leading to an improved fit of the spatio-temporal structure of on-going MEG data across all frequency bands.

## 2 Methods

### 2.1 Participants

The scanning of healthy participants was approved by the internal research board at the Center of Functionally Integrative Neuroscience (CFIN), Aarhus University, Denmark. The study was performed in accordance with the Declaration of Helsinki ethical principles for medical research and ethics approval was granted by the Research Ethics Committee of the Central Denmark Region (De Videnskabsetiske Komitéer for Region Midtjylland) (Ref 1-1072-252-13).

Sixteen participants with an average age of 24.75 (SD 2.54; 11 male, 5 female) were recruited through the online recruitment system at Aarhus University. Before inclusion into the study all participants received written and oral information about the entire study. All participants signed informed consent after receiving the information again on the first visit before any further screening and testing was completed. Participants were excluded if they suffered from psychiatric or neurological disorders or symptoms or had a history thereof. All participants were scanned on two separate days where the MEG data was acquired on the first day and the DTI data was acquired during the second visit.

### 2.2 MEG data collection and pre-processing

Empirical resting-state MEG (rs-MEG) data were acquired using a 306 channel Elekta Neuromag TRIUX system (Elekta Neuromag, Helsinki, Finland) located in a magnetically shielded room at the CFIN at Aarhus University Hospital, Denmark. All data were recorded at a sampling rate of 1000 Hz with an analogue filtering of 0.1–330 Hz. Approximately five minutes of resting-state was collected for each participant.

Before data collection, a three-dimensional digitizer (Polhemus Fastrak, Colchester, VT, USA) was used to record the participant's head shape relative to the position of four headcoils, with respect to three anatomical landmarks, which could be registered on the MRI scan (the nasion, and the left and right preauricular points). A structural MRI scan for each participant was acquired during a separate session during which DTI data were collected as well. The position of the headcoils was tracked during the entire recording using continuous head position identification (cHPI), providing information on the exact head position within the MEG scanner. This allows for accurate movement correction at a later stage during data analysis.

The raw MEG sensor data (204 planar gradiometers and 102 magnetometers) was downsampled from 1000 Hz to 250 Hz using MaxFilter and converted to SPM8 format. Using an in-house-built data viewer channels with excessive noise levels were marked and excluded from the following signal space separation. To attenuate interference originating outside of the scalp, signal space separation was applied to each data set using MaxFilter, applying its temporal extension as introduced in Taulu and Simola (2006). Data was once again converted to SPM8 format and remaining noisy segment and channels were excluded from further analysis, once again through visual inspection. In order to identify and remove signal originating from mains interference, ocular- and cardiac activity, as well as other non-neuronal interference each data set was then submitted to Blind Source Separation by temporal Independent Component Analysis (ICA), a tool that has been shown effective at separating artefactual signals from MEG sensor data (Mantini et al., 2011). Following Mantini et al. (2011) we used the FastICA algorithm (Hyvarinen and Oja, 2000, Vigario et al., 2000) as implemented in the FastICA toolbox for MATLAB (http://research.ics.aalto.fi/ica/fastica/). Each independent component (IC) was represented by an independent time-course and a mixing matrix describing each of the MEG sensors contribution to the given IC. Prior to ICA gradiometers and magnetometers were normalised by their respective minimum eigenvalues. The dimensionality of the ICA was set to 62. We used the following 4 criteria to identify artefactual ICs: i) if the spectrogram of the IC had a global maximum at 50 Hz and was of low kurtosis (<0.4) the IC was flagged as mains interference, ii) if the squared of the IC time course and the concomitantly acquired EOG correlated by more than 0.15, the IC was flagged as ocular interference, iii) if the squared of the IC time course and the concomitantly acquired ECG correlated by more than 0.15, the IC was flagged as cardiac interference, and iv) if the kurtosis of the IC time course was higher than 20 the IC was flagged as non-neural activity. This resulted in an average of ~ 3 rejected ICs across all data sets. Specifically, the artefactual ICs were removed by subtracting the contribution and the sensor data recomposed into its original channel dimensionality by the use of the SPM8 function *spm_eeg_montage,* where the montage-matrix, *M,* was estimated for each data set like the following:

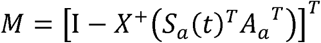

where I is the identity matrix of dimensionality [*number of channels* x *number of samples*]*, X^+^* is the pseudoinverse of the data matrix *X* [*number of channels* x *number of samples*] computed using an order of 62, *S_a_*(*t*) represents the time courses of the artefactual ICs, and *A_a_* [*number of artefactual ICs* x *number of samples*] their corresponding mixing matrices [*number of channels* x *number of artefactual ICs*]. Superscript *T* denotes the matrix transpose. The resulting de-noised sensor data was then used for further processing.

### 2.3 MEG data in Source Space

A scalar implementation of the LCMV beamformer was applied to estimate the source level activity of the MEG sensor data at each brain area (Van Veen et al., 1997, Sekihara et al., 2001, Woolrich et al., 2011). We used the widely-used and freely available AAL template (Tzourio-Mazoyer et al., 2002) to parcellate the brain into 90 non-cerebellar regions in line with previous studies of resting-state MEG data (Cabral et al., 2014b, Hindriks et al., 2015, Brookes et al., 2016). Note however that, although this anatomy-derived parcellation serves for the purpose of the current work, it is unlikely to perform as well as a function-derived parcellation involving PCA (Baker et al., 2014, Colclough et al., 2015).

For each individual subject, a structural T1 MRI scan was collected for the purpose of co-registration of the MEG data and the AAL parcellation template. The acquisition parameters for the T1 scan were: voxel size = 1 mm3; reconstructed matrix size 256×256; echo time (TE) of 3.8 ms and repetition time (TR) of 2300 ms. Each individual T1-weighted MR scans was co-registered to the standard MNI template brain through an affine transformation and further referenced to the space of the MEG sensors by use of the Polhemus head shape data and the three fiducial points. An overlapping-spheres forward model was computed, representing the MNI-co-registered anatomy as a simplified geometric model using a basis set of spherical harmonic volumes (Huang et al., 1999). One of the advantages of beamformers over other source-reconstruction strategies is that it reconstructs sources at any given location, independent from sources at other locations. Since we were interested in locating source activity at the AAL brain areas, we directed the beamformer to the center-of-gravity coordinates of the 90 non-cerebellar AAL areas as in Cabral et al. (2014b). The beamformer was applied to the sensor data band-pass filtered between 2-40 Hz, where the two sensor modalities (magnetometers and planar gradiometers) were combined by normalising them by the mean of their respective eigenvalues. The estimation of the data covariance matrix, necessary for the beamformer reconstruction, was regularised using the top 62 (minus number of rejected ICs) principal components. This yielded, for each subject, a [90 x *number of samples*] source-space data matrix, representing the spontaneous activity at the 90 AAL areas in the 2-40 Hz frequency range.

### 2.4 DTI data collection and Structural Connectivity

Two DTI datasets were acquired for each subject with opposite phase encoding directions. For the DTI data acquisition the following parameters were used: TR = 9000 ms, TE = 84 ms, flip angle = 90°, reconstructed matrix size of 106×106, voxel size of 1.98×1.98 mm with slice thickness of 2 mm and a bandwidth of 1745 Hz/Px. Furthermore, the data were collected with 62 optimal nonlinear diffusion gradient directions at b=1500 s/mm2. Approximately one nondiffusion weighted image (b=0) was acquired for every 10 diffusion-weighted images. Additionally, for the first acquisition we used an anterior to posterior phase encoding direction and the second acquisition was performed in the opposite direction.

The construction of the structural connectivity matrix (SC) followed the methodology from Cabral et al. (2012). Briefly, the regions defined using the AAL template (Tzourio-Mazoyer et al., 2002) were warped from MNI space to the diffusion MRI native space using the FLIRT tool from the FSL toolbox (www.fmrib.ox.ac.uk/fsl, FMRIB, Oxford) (Collins et al., 1994). Secondly, the connections between regions were estimated using probabilistic tractography with default parameters of the FSL diffusion toolbox (Fdt). The local probability distribution of fibre direction at each voxel was estimated following Behrens et al. (2003) and the probtrackx tool in Fdt was used for the automatic estimation of crossing fibres within each voxel (Behrens et al., 2007).

The connectivity probability from a seed voxel *i* to another voxel *j* was defined as the proportion of fibres passing through voxel *i* that reach voxel *j* using a sampling of 5000 streamlines per voxel (Behrens et al., 2007). This was extended from the voxel level to the region level, such that in an AAL region *i*, the connectivity probability P_*ij*_ from region *i* to region *j* is calculated as the proportion of sampled fibres in all voxels in region *i* that reach any voxel in region *j*. For each brain region, the connectivity probability to each of the other 89 regions within the AAL was calculated using in-house Perl scripts. As directionality of connections cannot be determined based on diffusion MRI, the undirected connectivity probability between regions *i* and *j* was defined as the average between P_*ij*_ and P_*ji*_. This undirected connectivity was considered as a measure of the structural connectivity C_*ij*_=C_*ji*_, resulting in a 90×90 symmetric weighted *matrix C* representing the network organization of each individual brain. A group averaged structural connectivity matrix <*C*> was obtained by averaging across all 16 healthy participants.

### 2.5 Whole-Brain Model

The whole-brain model consists of 90 coupled brain areas (nodes) defined according to the AAL parcellation scheme referred to above. We assumed that the local dynamics at the node level can be properly approximated to the normal form of a Hopf bifurcation (also known as Landau-Stuart Oscillator), in which all elements within a system (here, all neurons within a brain area) can be in an asynchronous noisy state below a given threshold, *a,* or display coherent oscillations above that threshold (see Figure 1 for an illustration). This is the canonical model for studying the transition from noisy to oscillatory dynamics from applied bifurcation theory (Kuznetsov, 1998). Recently, Freyer et al. (2011,2012) have nicely shown the usefulness, richness and generality of this type of model for describing EEG dynamics at the local node level. Here, we focus on how those local noisy oscillators interact through the anatomical network to generate spontaneous spatio-temporally structured fluctuations. Within this model, each node *j* of the network was modelled by a normal Hopf bifurcation with a fundamental frequency *f_f_=ω/2π*equal for all nodes as:

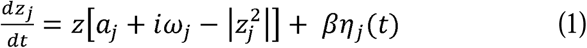

where

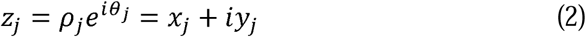

**Figure 1.**
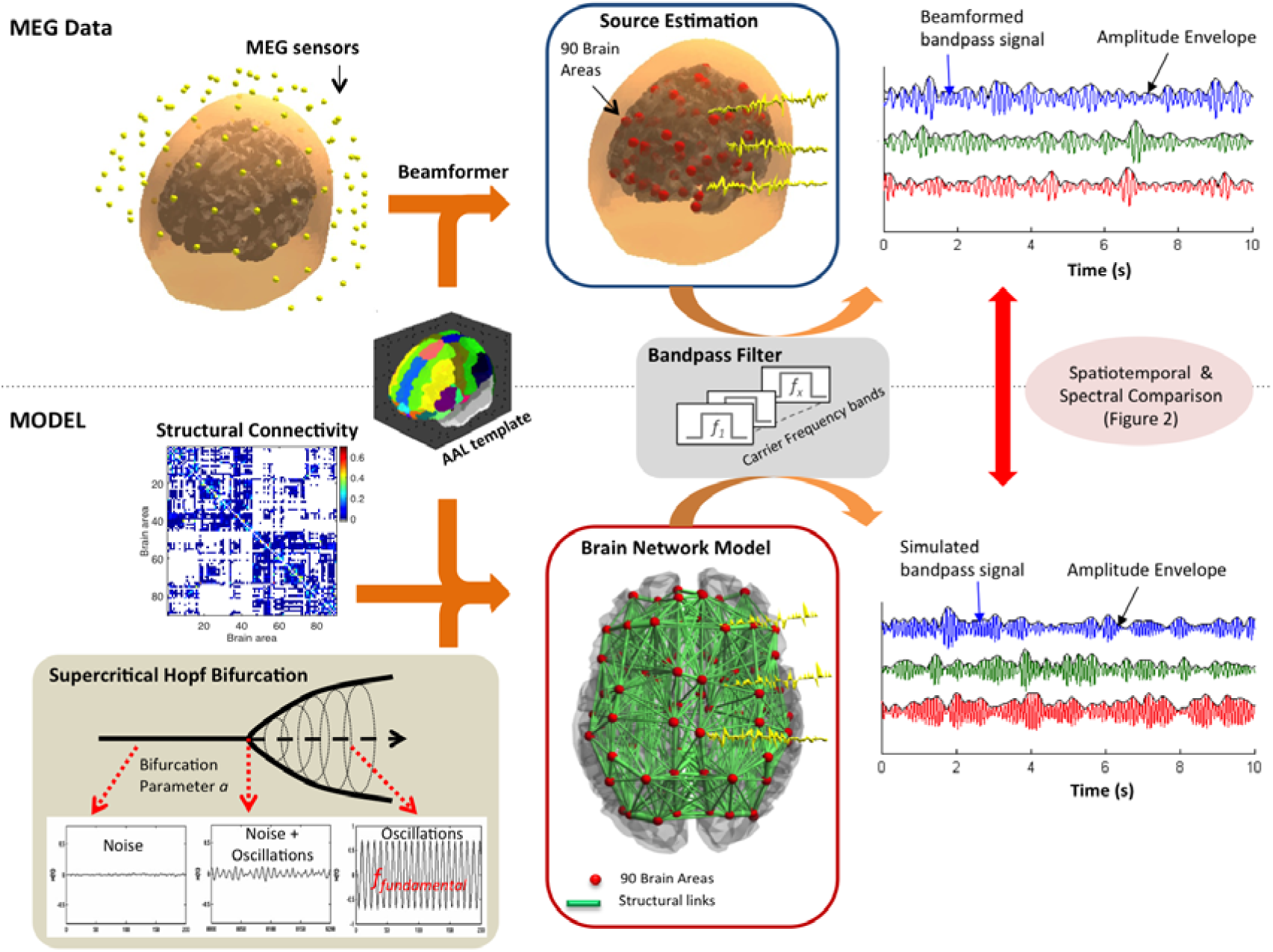
Data processing pipeline. MEG data. The broad-band MEG sensor signals are beamformed into source space to the centre-of-gravity locations of 90 brain areas defined in the AAL template. The signal is bandpass-filtered in narrow carrier frequency bands and the corresponding amplitude envelopes are computed using the Hilbert transform. **Model:** Structural connectivity (SC) between the 90 AAL brain areas is computed from DTI-based tractography. Each entry SC(n,p) contains the average proportion of sampled fibres in region *n* that reach region *p*. The local dynamics of each brain area is modelled using the normal form of a Hopf bifurcation at the critical point between a quiescent state a (represented by noise) and the emergence of oscillations (which occur at a predefined fundamental frequency). The simulated data is subsequently bandpass-filtered in narrow carrier frequency bands and the corresponding envelopes are computed in the same way as the empirical MEG data.

Where *η_j_*(*t*) is additive Gaussian noise with standard deviation ***β***. Inserting equation (2) in equation (1) and separating the real part in equation (3) and the imaginary part in equation (4) we obtain:

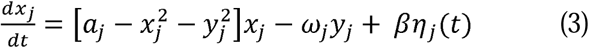

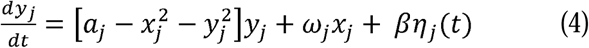

The global dynamics of the whole-brain model results from the mutual interactions of local nodes coupled through the underlying anatomical connectivity. The structural matrix *C_ij_* denotes the density of fibres between cortical areas *i* and *j* as extracted from the DTI-based tractography, averaged across the 10 subjects. Thus, the whole-brain dynamics was defined by the following set of coupled equations:

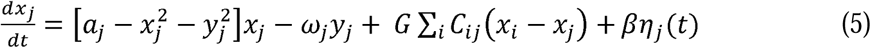

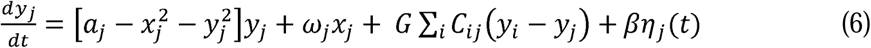

where *X_j_* is the simulated MEG signal of each node *j*. Each node *j* has a supercritical bifurcation at *a_j_=0,* such that for *a_j_<0* the node is in a stable fixed point at *(X_j_,y_j_)=(0,0)* and is represented by neuronal noise (corresponding to the asynchronous firing of neurons), whereas for *a_j_>0* the system switches to a pure oscillatory state (corresponding to the synchronized firing of neurons at a fundamental frequency *f_f_ =ω/2π*). When the system is operating near the bifurcation (*a=0*) in the presence of noise, noisy fluctuations may induce temporary excursions to the oscillatory regime. During such excursions, the local node will display the temporary emergence of oscillations at the fundamental frequency *f_f_*. Importantly, regions receiving correlated input are more prone to generate oscillations simultaneously, leading to correlated envelopes of carrier oscillations around *f_f_*. Since we are specifically interested in investigating such type of mechanism, we considered all nodes to be operating in the critical regime, i.e. *a_j_=0* for all nodes. *G* is a global scaling factor (global conductivity parameter scaling equally all synaptic connections) and serves as the control parameter to find the optimal dynamical working region where the simulations maximally fit the empirical data. In order to fall in the same range of *G* as in previous studies (Deco et al., 2016), the maximum value of *C_ij_* was set to 1 and multiplied by a scaling factor of α=0.2 as C=α*C/max(C(:)). This procedure only rescales the arbitrary units of the coupling strength *G* (which we varied between 0 and 1) and of the standard deviation of additive Gaussian noise (which was set to ***β***=0.02) without affecting the overall model results.

Regarding the fundamental frequency at which brain areas oscillate, we explored a range of frequencies between 1 and 30Hz. The selection of frequencies was purely phenomenological, with the intention of verifying if Hopf bifurcation models could reproduce the envelope correlations detected in resting-state MEG studies, which are maximal for carrier oscillations between 8 and 20Hz (Brookes et al., 2011b, Hipp et al., 2012). Actually, it is known that neural masses mostly resonate in the gamma-frequency range (>30Hz), whereas slower cortical rhythms (<30Hz) are currently believed to arise from thalamo-cortical interactions (Buzsaki, 2006, Palva and Palva, 2007, Engel et al., 2013). Irrespective of their genesis, empirical evidence shows that the latter emerge spontaneously in the cortex and in a correlated fashion during rest. Following this evidence, we model their emergence as a transition to the oscillatory state with a Hopf bifurcation model.

For the single-frequency model, simulations were performed for a range of fundamental frequencies *f_f_*=1:2:29Hz with *f_f_* equal for all brain areas. Simulations were subsequently run with the selected optimal frequency *f_f_*=12Hz. In order to simulate a multi-frequency model, we computed simulations with *f_f_*=4,8,12,16,20,24,28 Hz (as in the single-frequency case), with homogeneous coupling G for all fundamental frequencies, and analysed each simulation in the corresponding carrier frequency band (see Figure 3). All simulations were run for a duration of 3200 seconds, which corresponds to the total duration of the MEG recordings from the 16 subjects. Since time delays do not contribute for this specific mechanism and the envelope fluctuations are intrinsically slow, we have assumed instantaneous transmission between nodes to reduce the complexity of the simulations.

### 2.6 Frequency-specific Amplitude Envelope

Following recent developments in the analysis of spontaneous MEG data, we focused our analysis on the amplitude envelopes of distinct carrier oscillations, considering the narrowbands [*f_carrier_-2, f_carrier_+2* Hz] with *f_carrier_=4:2:28* Hz) (see Figure 1) (Brookes et al., 2011a, Cabral et al., 2014b). Note that the power of an oscillation is proportional to its squared amplitude, so this is comparable to using the band-limited power (BLP) of the MEG signals. To obtain the amplitude envelopes, first the source-reconstructed MEG signals (or simulated data) were band-pass filtered within the narrowbands [*f*-2,*f*+2 Hz] and the instantaneous amplitude of each narrowband signal was calculated using the Hilbert transform. The Hilbert transform represents a narrowband signal, *s*(*t*), in the time domain as a rotating vector with an instantaneous phase, *φ*(*t),* and an instantaneous amplitude, *A(t)* such that *s*(*t*) = *A*(*t*)cos (*φ*(*t*)). Further, we consider solely the ultra-slow fluctuations of the amplitude envelope *A*(*t*) by low-pass filtering at 0.2 Hz as this is known to maximize meaningful resting-state FC in MEG studies (Brookes et al., 2011a, Hipp et al., 2012). Indeed, Hipp and colleagues (2012) have shown that, the slower the envelope component, the higher the envelope correlation between distant, yet functionally related, brain areas. This slow component of the envelopes of each brain node at a given carrier frequency is the signal used in subsequent analysis of temporal correlations and phase dynamics (Figure 2).

**Figure 2.**
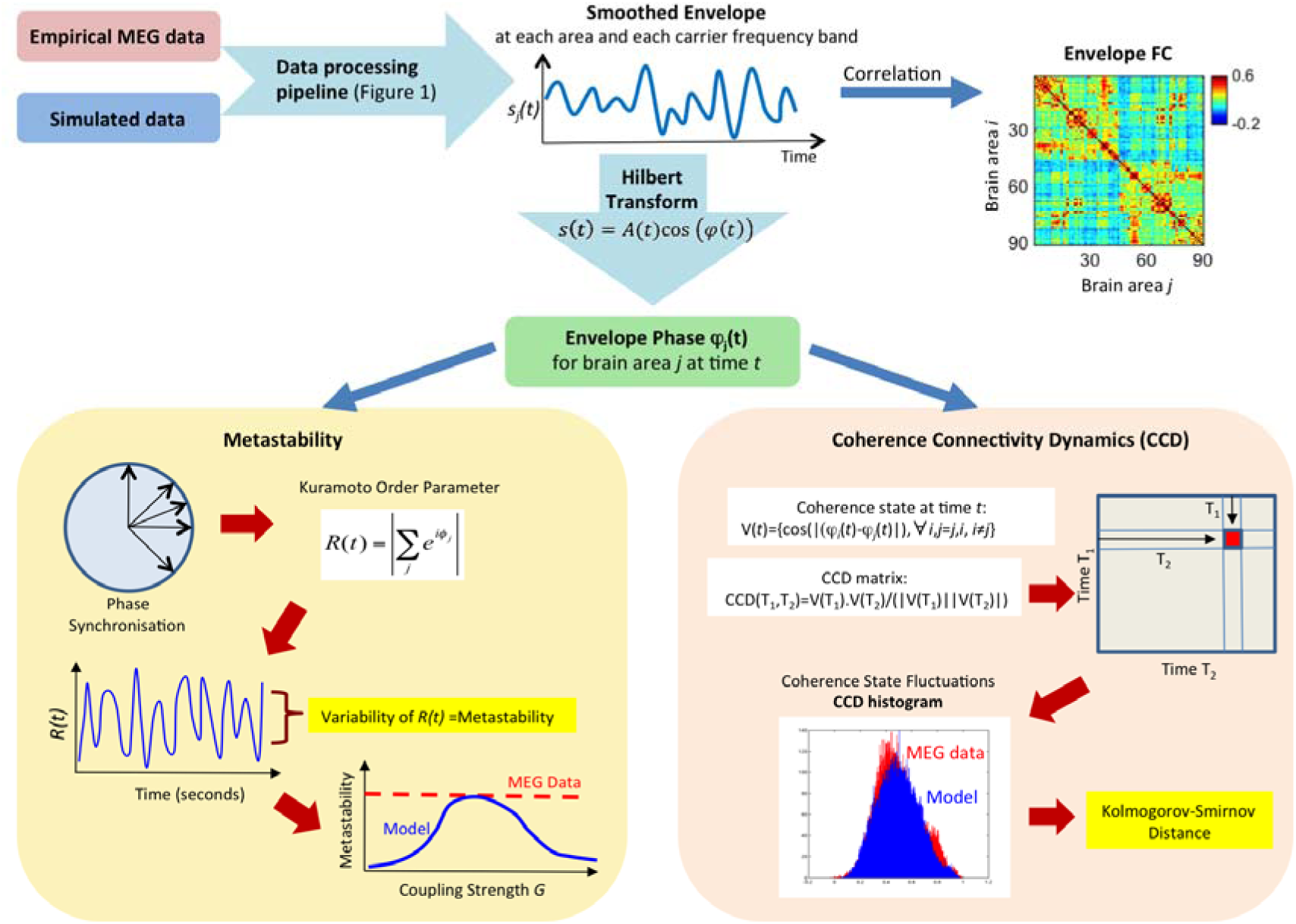
Data analysis pipeline. **Top -** The analysis is performed separately for each frequency band, starting with the low-pass filtered (<0.2Hz) envelopes of both pre-processed and source-reconstructed MEG signals and simulated model data. The **Envelope FC** is the Pearson’s correlation between the low-pass filtered envelopes from all pairs of brain areas. For the phase analysis, we apply the Hilbert transform to the envelope and obtain the Envelope Phase at each instant of time. **Bottom left panel -** Instantaneous phase synchronization *R(t)* is calculated using the Kuramoto Order Parameter and its variability is used as a measure of the system’s **Metastability**. The dynamical working point of the model is adjusted in order to approximate the metastability observed in resting-state MEG data. **Bottom right panel -** To obtain the matrix of **Coherence Connectivity Dynamics** (**CCD**) we first calculate the Coherence state V(t), which is a vector with the instantaneous phase coherence between all pairs of nodes. The cosine similarity between V(t_1_) and V(t_2_) is reported in CCD(t_1_,t_2_), providing a picture of the evolution of coherence states over time. The distribution of values in the upper triangular part of CCD the matrix is compared between empirical and model experiments using the Kolmogorov-Smirnov Distance, in order to search for the optimal model parameters that generate a Phase coherence dynamics similar to the one observed in MEG data from healthy subjects at rest.

### 2.7 Envelope FC

The Envelope FC was defined as the matrix of Pearson’s correlations of the slow components of the amplitude envelopes of MEG signals at a given carrier frequency over the whole acquisition time (see Figure 2). For each carrier frequency band, the Envelope FC matrices were averaged across participants, in line with previous resting-state Envelope FC studies (Cabral et al., 2014b, Brookes et al., 2016).

### 2.8 Metastability

In order to analyse the envelope connectivity dynamics in the phase domain, we first calculated the instantaneous phase of the envelopes of each brain area at each carrier frequency band using the Hilbert transform (see Figure 2) (Glerean et al., 2012). We refer to metastability as a measure of the variability of phase configurations as a function of time, i.e. how the global synchronization between brain areas fluctuates across time (Shanahan, 2010, Wildie and Shanahan, 2012, Cabral et al., 2014b). Thus, we measured the metastability as the standard deviation of the Kuramoto order parameter, which is defined by the following equation:

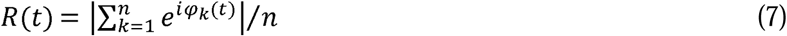

where φ_*k*_(*t*) is the instantaneous phase of the narrowband envelope at node *k.* The Kuramoto order parameter measures the global level of synchronization of the ensemble of *n* oscillating signals. Under complete independence, the *n* phases are uniformly distributed and thus *R* is nearly zero, whereas *R* = 1 if all phases are equal (full synchronization). We computed the metastability of the empirical MEG and simulated signals at each carrier frequency using the Hilbert-derived phases of the slow envelope fluctuations.

### 2.9 Coherence Connectivity Dynamics (CCD)

In order to characterize the time-dependency of the envelope correlations, we estimated a frequency-specific version of the FCD matrix (Hansen et al., 2015) that we call Coherence Connectivity Dynamics (CCD) (see Figure 2). The CCD is constructed following these steps: 1) Calculate the instantaneous phase of the envelopes of each brain area at each carrier frequency band; 2) Compute a vector V(t) representing the instantaneous coherence state, containing the cosine of the absolute phase difference between all undirected pairs of nodes; 3) Calculate the CCD which is a time-versus-time symmetric matrix where the (t_1_, t_2_) entry is defined by the cosine similarity (i.e. normalised vector product) between the instantaneous coherence vectors at times t_1_ and t_2_. Epochs of stable coherence states are reflected as blocks around the CCD diagonal. The histogram of CCD values contains valuable information about the time-dependencies of envelope phase dynamics in empirical data and was used as the most sensitive measure to fit with the model. At each carrier frequency band, the upper triangular elements of the empirical CCD matrices (over all participants and sessions) were computed and their distribution was compared with the distribution of simulated CCD values (for each value of *G*) using the Kolmogorov-Smirnov distance (KS-distance), which quantifies the maximal difference between the two cumulative distribution functions (see Figure 2). As such, the KS-distance serves as a fitting function to minimize the differences between the real and simulated CCD histograms.

## 3 Results

In order to emulate the global characteristics of spontaneous whole-brain dynamics observed in empirical MEG data from a group of healthy humans we used a whole-brain model linking the underlying anatomical structural connectivity (derived from DTI tractography) with the local dynamics of each brain area. The spatial and temporal structures of spontaneous MEG fluctuations were characterized focusing on three observables: 1) the Envelope FC; 2) the Coherence Connectivity Dynamics (CCD); and 3) the metastability index. The Envelope FC describes the Pearson’s correlations of the envelopes between different brain areas over the entire recording time. The CCD describes the temporal evolution of envelope coherence states. The metastability characterizes the richness of the dynamical repertoire by means of the variability of the global level of synchronization between envelopes (see *Methods* for details). These three measures were calculated for each carrier frequency considered in this study: *f_carrier_=*4:2:28 Hz in windows [*f_carrier_-*2 Hz, *f_carrier_+*2 Hz]. We performed the same analysis on both empirical and model data. Figure 1 sketches the general processing pipeline. Figure 2 describes the analysis pipeline for computing the three measures, namely the Envelope FC, the CCD and the metastability.

### 3.1 Single-frequency model

As a first step, we started by evaluating the performance of the classical model, considering only one fundamental oscillation frequency at a time, identical for all brain areas, varying *f_f_* between 1 and 29 Hz with increments of 2Hz (See Figure 3A). For each fundamental frequency considered, we analysed the resulting simulations in all carrier frequency bands. In Figure 4A we show the mean FC and the metastability of empirical resting-state MEG activity as a function of the carrier frequency band [*f*-2 Hz, *f+2* Hz]. We find that correlations are stronger and the level of synchrony fluctuates more for envelopes of carrier oscillations between 10 and 16Hz. That particular range of carrier frequencies was already found to maximize inter-area envelope correlations in a number of MEG resting-state studies (Brookes et al., 2011a, Hipp et al., 2012, Cabral et al., 2014b), and we extend those results here for the second order statistics, i.e. for the degree of richness in the exploration of the dynamical repertoire given by the metastability index.

**Figure 3.**
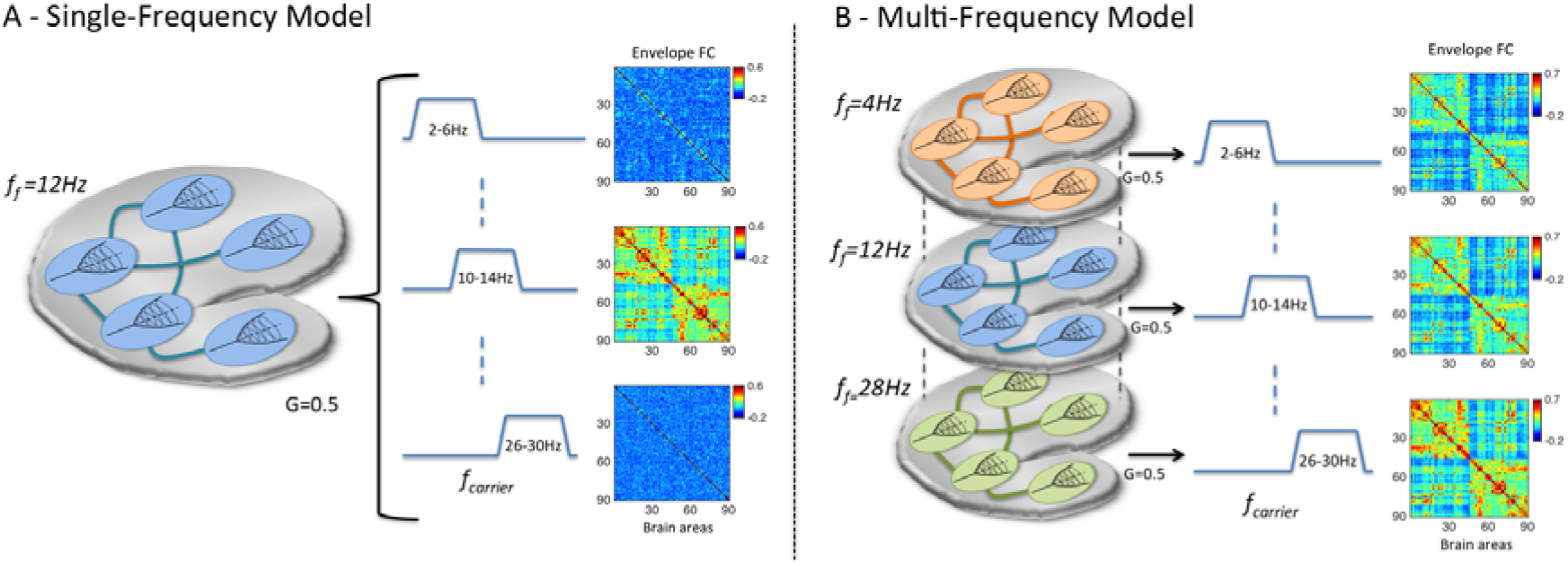
Illustration of the two different modelling approaches considered. A. In the single-frequency model, only one fundamental oscillation frequency **(*f_f_*),** identical for all 90 brain areas, is considered at a time. The output is filtered in narrow carrier frequency bands **(*f***_*carrier*_) and compared with empirical data. **B** – In the multi-frequency model, each brain area generates rhythms in different fundamental frequencies. Whole-brain simulations are performed independently for each fundamental frequency (as in the single-frequency model). The simulated signals at each fundamental frequency are then filtered only in the corresponding carrier frequency band. As such, the Envelope FC matrices from each carrier band are originated by a different set of Hopf-bifurcation models located in each brain area.

**Figure 4.**
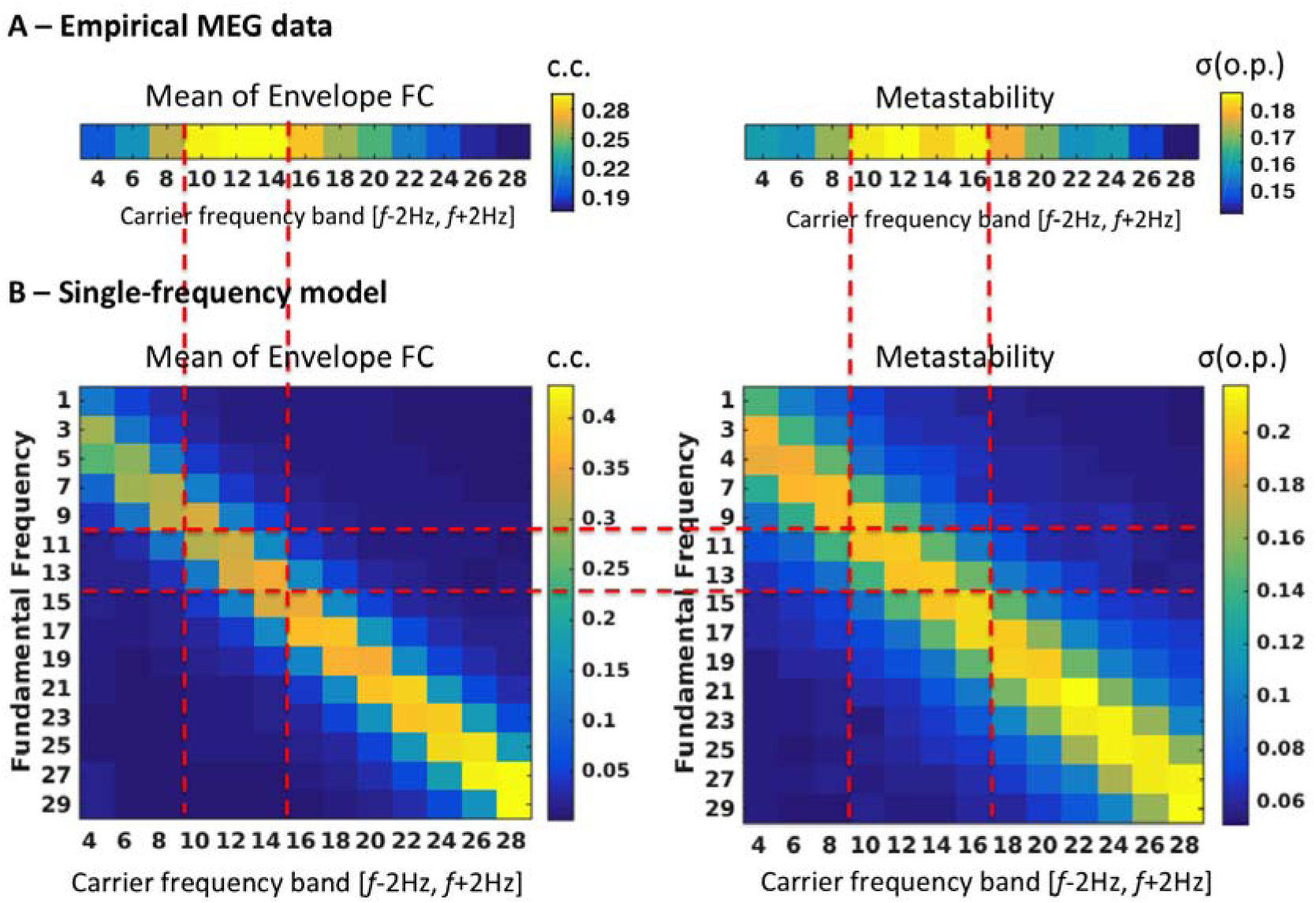
Finding the optimal fundamental frequency for the single-frequency model. **A** Properties from resting-state MEG data that we aim to reproduce with the model: (Left) Mean value of the envelope FC matrices at each carrier frequency (c.c. = correlation coefficient). The envelope correlations are stronger for 10-14Hz carrier oscillations. (Right) The metastability - measured as the standard deviation of the order parameter, σ(o.p), of envelope phases - has a broad peak between 10 and 16Hz. B – Results obtained from the single-frequency model with a range of fundamental frequencies (y-axis) to compare with the empirical data. The red dashed-lines serve as visual guidelines to match the peaks from panel A with the simulated values in panel B. We find that, using a fundamental frequency between 11 and 13Hz, the simulated signals show a peak in mean envelope FC (Left) and metastability (Right) for same range of carrier frequencies observed empirically. As such, *f_f_*=12Hz was selected as the optimal fundamental frequency for the single-frequency model.

In Figure 4B, we calculate the same measures but for each fundamental frequency assumed in the classical Hopf model (y-axis) as a function of the carrier frequencies (x-axis). The global coupling of the model was initially set to G=0.5, which falls in the optimal range where the simulations best replicate the empirical measures, as we will show in Figure 5. From the comparison between simulations and experimental results, we find that the most suitable fundamental frequency lies between 11 and 13Hz. Setting oscillations in the model in that frequency range, the mean FC and the metastability peak between 10-14Hz, quantitatively reproducing the respective values observed in the empirical MEG data.

**Figure 5.**
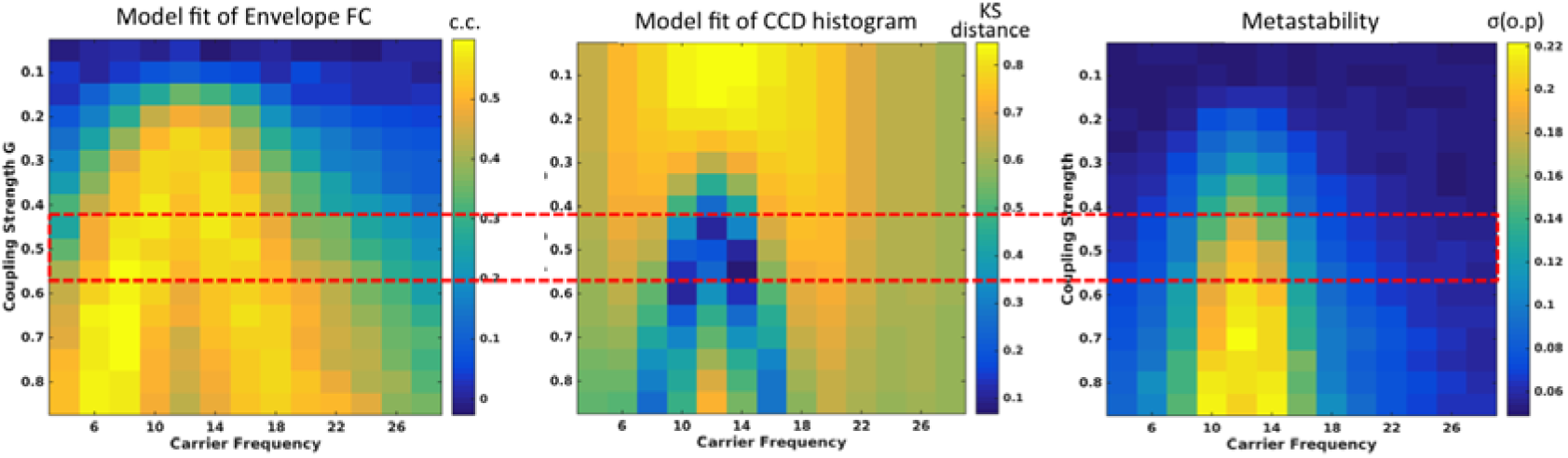
Optimizing the global coupling strength G for the single-frequency model with *f_f_*=12Hz. (Left) Correlation **coefficient (c.c.)** between the empirical and simulated Envelope FC matrices at each carrier frequency as a function of the global coupling strength G. (**Middle**) Kolmogorov-Smirnov (KS) distance between the empirical and simulated CCD histograms at each carrier frequency as a function of the global coupling G. (**Right**) Index of metastability (measured as the standard deviation of the order parameter, σ(o.p), of the simulated signals at each carrier frequency, as a function of the global coupling G. The red dashed-lines serve as visual guides to highlight the range of optimal coupling strength.

Selecting an optimal fundamental frequency of 12 Hz, we then explored the level of fitting of the three measurements: FC, CCD and metastability, in all carrier frequencies, as a function of the global coupling parameter *G* of the model (Figure 5). To define the optimal coupling value, we considered the 3 observables but focused mainly in minimizing the Kolmogorov-Smirnov distance between the empirical and simulated CCD histograms, which capture the time-dependencies of the envelope fluctuations, a feature of high sensitivity in the model. Actually, the correlation between empirical and simulated FC matrices, which is a traditional fitting measure used in most resting-state models, does not capture the temporal dimension and can be high for a broad range of G. Moreover, the metastability only captures the intensity of coherence fluctuations without informing about the time-scales at which these occur. As such, we chose to optimize G such that the temporal dynamics in the model best fits the empirical data in the range between 8 and 16Hz, where resting-state MEG has shown more meaningful envelope FC. As can be seen in Figure 5, this occurs for 0.4<G<0.6, so we chose G=0.5.

An example of the simulated data with *f_f_* = 12 Hz and *G*=0.5 is shown in Figure 6 for two representative brain areas. When the simulated signals are bandpass-filtered between 10-14Hz, we find that the amplitude envelope fluctuates over time. At a global level, we find that the mean amplitude of the bandpass-filtered signals over all brain areas correlates strongly with the zero-lag phase synchrony of the underlying carrier oscillations (cc=0.77), as shown in Figure 6C.

**Figure 6.**
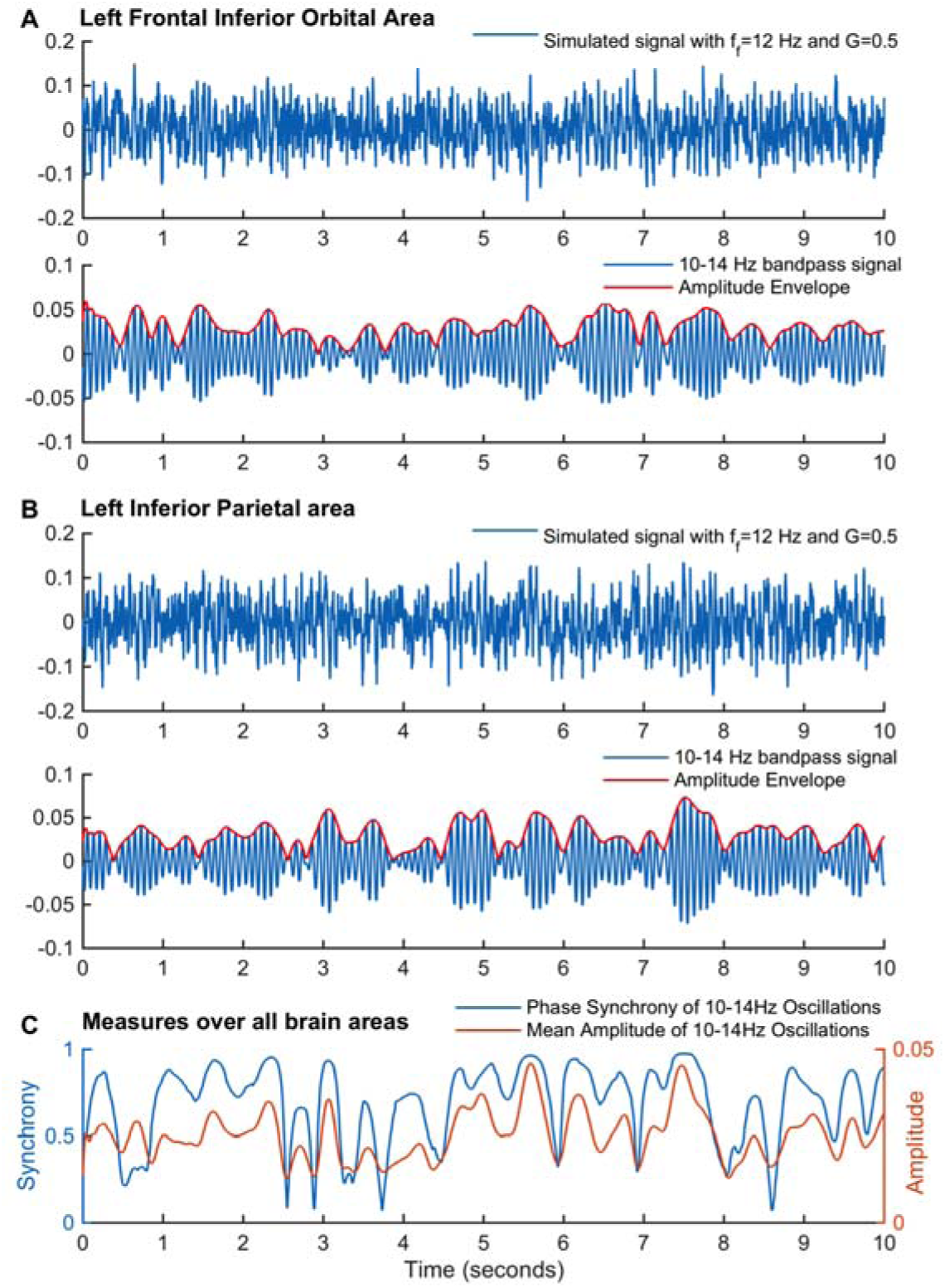
Simulated signal in two brain areas before and after band-pass filtering. A and B – (Top) Simulated signal obtained with the Hopf model with *f_f_* = 12 Hz and *G*=0.5. (Bottom) The same signal bandpass filtered between 10 and 14 Hz (blue) with the corresponding amplitude envelope obtained using the Hilbert transform (red). Plots are shown for 10 seconds of simulations and for two randomly selected seeds, the left Frontal Inferior Orbital area (A) and the left Inferior Parietal area (B). C – The phase of the 10-14Hz band-pass filtered signals was obtained for each brain area using the Hilbert transform and the global synchrony degree was estimated using the Kuramoto Order Parameter. Overall, the synchrony degree (blue) correlates strongly with the mean amplitude (red) of 10-14 Hz oscillations (cc=0.77).

The performance of the single-frequency model with fundamental frequency *f_f_=* 12 Hz and coupling strength G=0.5 is shown in Figure 7 in comparison with empirical MEG data. The panel (A) plots the correlation between the empirical and simulated Envelope FCs and the Kolmogorov-Smirnov distance between the empirical and simulated CCD histograms, both as a function of the carrier frequency. The single-frequency Hopf model is clearly able to achieve a good fit for the limited -but most relevant (see Figure 4A)- range of carrier frequencies between 10 and 14Hz. The same happens with the metastability (Figure 7B). However, the performance of the model decreases as the difference between the fundamental frequency and the carrier frequency increases. In order to visualize the similarity between simulations (with *f_f_*=12Hz, *G*=0.5) and empirical resting-state data for the carrier band centred at *f_carrier_*=12Hz, we report in Figure 7C the corresponding Envelope FC matrices, the CCD matrices, CCD distributions and the evolution of the Envelope Synchronization over time (defined in equation 7). Both empirical and simulated CCD matrices show a checkerboard pattern, which is indicative of time-dependencies, i.e, the coherence between envelopes (CC) is not constant over time but instead alternates between periods of increased temporal similarities (lasting up to several seconds) with periods of incoherence. Similar checkerboard patterns have been reported by Hansen et al. (2015) in FCD matrices, where the correlation matrices of BOLD signals were compared over sliding-windows. The evidence of temporal dependencies is complemented by the long-tailed shape of the CCD distributions. If the patterns of CC were constant over time, all CCD values would be shifted to 1. On the other hand, if CC states were unrelated over time, the CCD distributions would be shifted to zero. Instead, we find a peak at relatively low values (corresponding to the periods of weak temporal similarities) together with a long tail towards high CCD values, indicating the existence of periods of high similarity between CC states. Moreover, as can be seen in the right plot, the synchrony degree of envelope phases fluctuates strongly over time (i.e. high metastability) meaning that envelopes synchronize and desynchronize over time. In the model (bottom), a similar temporal dynamics of envelope connectivity is achieved for *G*=0.5, where interactions between brain areas through the structural connectome in the presence of noise lead to temporary excursions to the oscillatory state. Constant transitions in and out of the oscillatory state induce correlated envelope fluctuations of 12Hz oscillations between structurally connected brain areas (as seen in the envelope FC matrix). Over time, we observe similar fluctuations in the synchrony degree as observed in the experimental data. However, although the single-frequency model is a good candidate to explain the dynamics occurring in the specific frequency range around the fundamental frequency (here for the carrier bands 8-12Hz, 10-14Hz and 12-16Hz), it fails to reproduce the dynamics when we consider slower or faster carrier oscillations, as can be seen by the increased distance between CCD histograms (A), the decreased Envelope FC fitting (A) and the lower metastability (B), indicating the absence of metastable coherent states in other frequency ranges.

**Figure 7.**
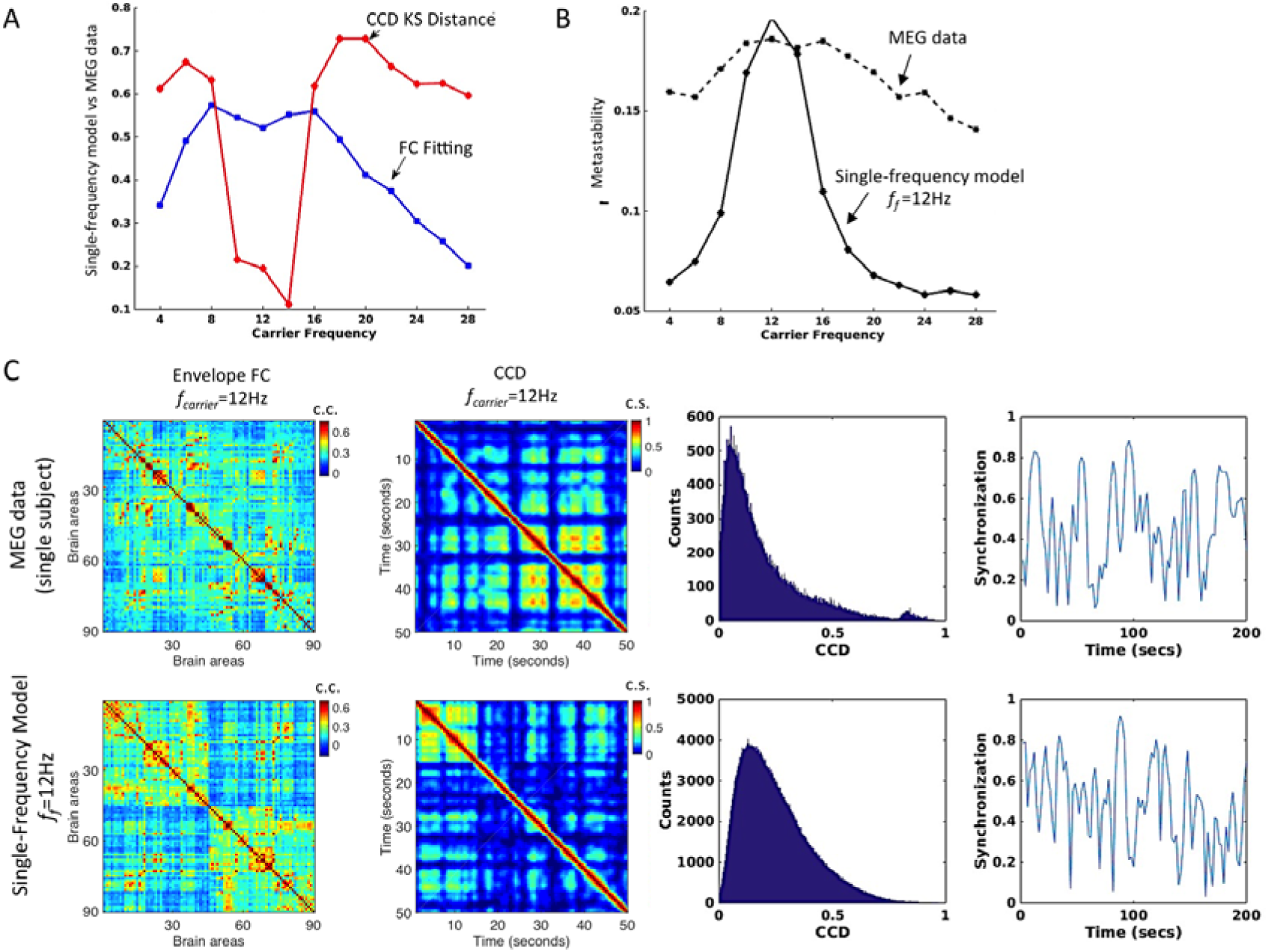
Performance of the single-frequency model with *f_f_*=12Hz and *G*=0.5. A – (Blue) Correlation between the envelope FC matrices (upper triangular part only) obtained with the single-frequency model and from resting-state MEG data as a function of the carrier frequency. (**Red**) Kolmogorov-Smirnov distance between the histograms of the values in the upper triangular part of the CCD matrices obtained from empirical data and with the model. **B** – Index of Metastability, measured as the coherence variability of envelope phases. **C** – Example from a single-subject MEG recording (top) and from simulations with the single-frequency model with *f_f_*=12Hz and *G*=0.5 (bottom) of the Envelope FC matrices (c.c = correlation coefficient), the CCD matrices distributions (c.s. = cosine similarity) and temporal evolution of the Envelope Phase Synchronization (Order Parameter) for the carrier frequency *f_carrier_=*12. The CCD matrices are shown only for 50 seconds for a better illustration of the *checkerboard effect,* indicative of time-dependencies, similarly to what is obtained in fMRI studies of Functional Connectivity Dynamics (Hansen et al., 2015).

### 3.2 Multi-frequency model

Considering that the brain operates at multiple frequency levels and has the capacity to generate oscillations in a broad range of frequencies within a brain area, a multi-frequency model may be formulated, as sketched in Figure 3 (right). In this framework, spontaneously generated oscillations interact in the structural network, giving rise to structured amplitude envelopes with fluctuating phase coherence (i.e. metastability) in all frequency ranges.

Previously, in Figure 4B, we observed that a shift in the fundamental frequency of the Hopf model led to an associated shift in the frequency at which the envelopes correlate the most and where the corresponding phase coherence fluctuated the most. Moreover, as shown in Figure 5, the scaling of the coupling strength G allows fine-tuning the amount of metastability and the associated dynamics of Phase coherence states (as captured by the CCD histograms). This suggests that a multi-Hopf model would consequently be able to describe the FC and metastability equally well in the whole range of carrier frequencies. In order to show this, we simulated a multi-frequency model using the fundamental frequencies 4, 8, 12, 16, 20, 24 and 28 Hz in parallel. For all fundamental frequencies, we considered the same value of global coupling *G=0.5.* In Figure 8A and B we show that now all the three fitting measures are similarly good across the whole range of carrier frequencies considered, namely the correlation between simulated and empirical Envelope FCs is kept high above 0.45, the Kolmogorov-Smirnov distance between CCD histograms is always shorter than 0.2 and the metastability profile is similar to the empirical one, with a peak at 12 Hz.

**Figure 8.**
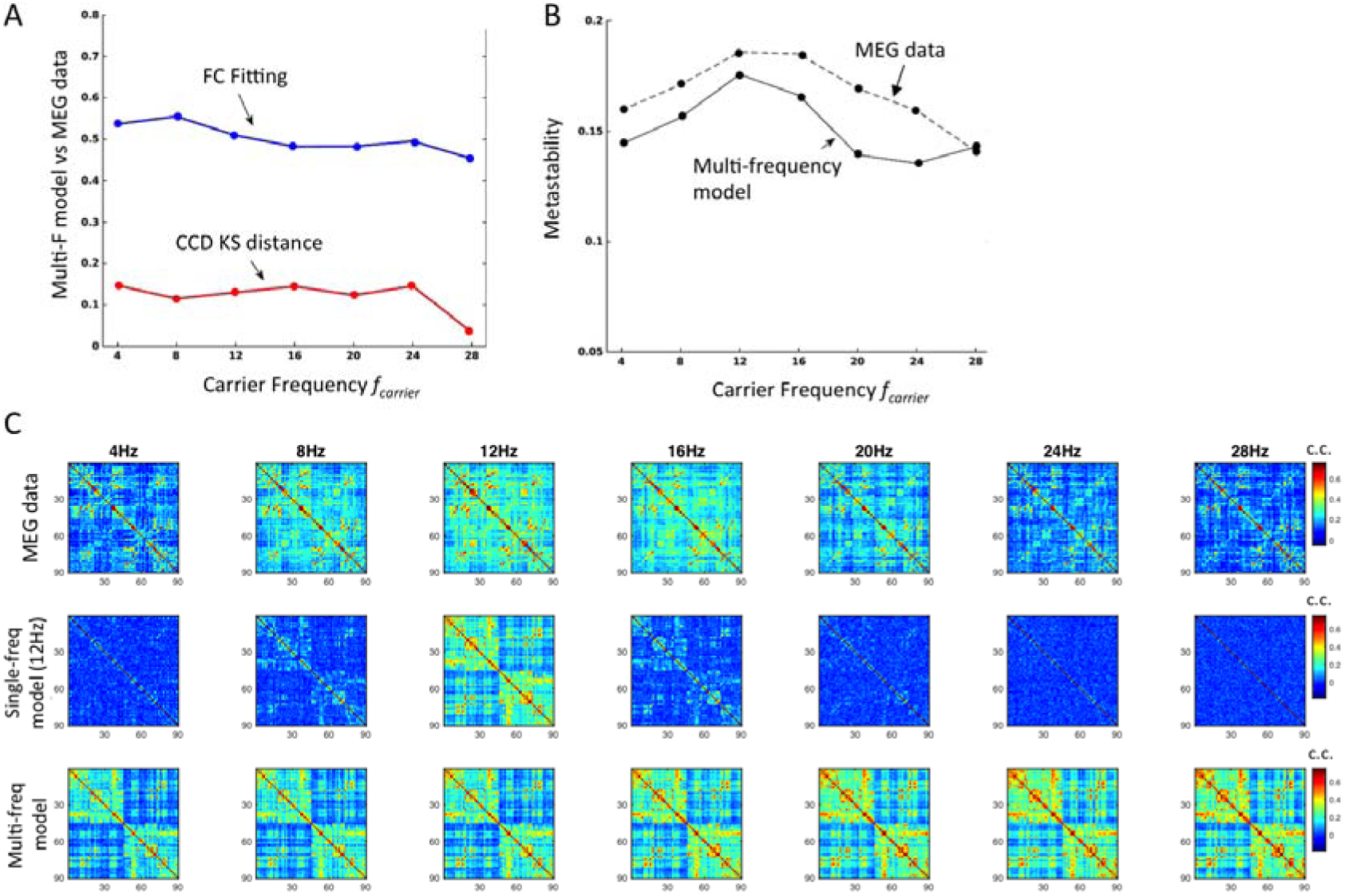
Performance of the multi-frequency model and comparison between Envelope FCs across modalities. **A** - Correlation between simulated and empirical Envelope FCs (Blue) is kept high above 0.45 across all carrier frequencies and the Kolmogorov-Smirnov (KS) distance between CCD histograms remains low (<0.2). **B -** The metastability profile is similar to the empirical one across the spectrum of carrier frequencies, with a peak around 12 Hz. **C** - Envelope FC matrices obtained from empirical MEG data (top) and from simulations with the single-frequency model (middle, *f_f_*=12Hz) and the multi-frequency model (bottom) for the range of non-overlapping carrier bands considered.

Finally, in Figure 8C, we show the empirical and simulated envelope FC matrices for the range of non-overlapping carrier bands considered. The empirical envelope FC matrices (top) are spatially structured across the whole range of carrier frequencies considered, with some pairs of brain areas displaying stronger envelope correlations than others. In the current model, the main source of correlated input driving the simultaneous emergence of oscillations is the structural connectivity matrix *C,* obtained from DTI-based tractography. Considering the single-frequency model with *f_f_* = 12Hz (middle) we find that the envelope FC matrix is shaped by *C* for the carrier band centred at the fundamental frequency of 12Hz, with some structure appearing in the neighbour bands around 8 and 16Hz. However, when the carrier frequency band is more distant from the fundamental frequency, the envelope FC matrices appear totally random and the correlation with empirical matrices decreases considerably (as shown in Figure 8A). On the other hand, considering the multi-frequency model, correlated envelope fluctuations appear in each frequency band resulting in structured envelope FC matrices across the frequency spectrum (bottom). Since in each frequency layer the nodes are coupled according to the same structural connectivity matrix *C* (which shapes the envelope correlations) the FC matrices are similar across the frequency spectrum, only varying the intensity of the correlations according to the global coupling *G*. In fact, for the current model to form frequency-specific functional networks, a different structural network should be considered at each frequency layer, with loss of biophysical realism. Additionally, if the structural connectivity was randomized, the FC matrices would consequently appear randomized and the correlation with the empirical FCs would drop drastically.

In the current model, and for a phenomenological proof of concept, we considered only two extreme scenarios. However, including a suitable number of optimized frequencies would not only increase the performance with regards to the single-frequency model but also make it more realistically plausible than the multi-frequency model. For instance, considering only two natural frequencies corresponding to typically observed alpha and beta peaks, most of the CCD variance and envelope FC structure would be replicated in the frequency range where maximal metastability is observed in real MEG recordings.

## 4 Discussion

Over the last decade, a number of whole-brain network models have been used to investigate the mechanisms underlying resting-state activity observed with fMRI, proposing different scenarios for the origin of structured BOLD-signal fluctuations. However, due to its intrinsic limitations, fMRI provides only a narrow picture of the complex dynamics governing brain activity at rest. Recent MEG observations have revealed meaningful envelope correlations of band-limited signals, imposing novel constraints to consider in whole-brain models of spontaneous activity.

Following the expanding trend focusing on the envelope dynamics of band-limited carrier oscillations, we extract the envelopes from resting-state MEG signals on multiple carrier frequency bands and analyse not only their correlation maps across the brain (Envelope FC) but also explore new grounds by considering the coherence of envelope phases over time (i.e. the metastability) and explore time-dependencies using a measure of coherence connectivity dynamics (CCD). Using a computational model, we test two mechanistic scenarios for the origin of spatio-temporally organized envelope fluctuations of narrow-band carrier oscillations observed in resting-state MEG signals: one scenario where brain areas only resonate at a single oscillatory frequency, and another where brain areas resonate in parallel at multiple different frequencies.

### 4.1 Empirical evidence from resting-state MEG

Looking at the MEG data from healthy subjects at rest, we find that the envelopes appear more strongly correlated for carrier waves between 10-16Hz, corroborating previous findings (Brookes et al., 2011a, Brookes et al., 2011b, Hipp et al., 2012, Cabral et al., 2014b). Moreover, we find that the coherence of envelope phases fluctuates strongly for the same range of carrier waves, which is indicative of metastability in the system. This reveals a rich dynamical regime, in which the envelope phases are neither fully synchronized nor completely incoherent over time, but fluctuate between states of partial synchronization that exist in the dynamical repertoire of the system. In addition, considering the CCD, we find that the patterns of envelope coherence replicate over time, with a long-tailed distribution of CCD values. These three observables are used to constrain the models in order to identify the optimal working point at which the brain is most likely to operate during rest.

### 4.2 Models and mechanisms of resting-state MEG FC

Despite the empirical evidence of frequency-specific envelope FC, the mechanism at the genesis of this phenomenon remains unclear. A successful model of resting-state MEG data must generate frequency-specific envelope FC, where a given pair of brain areas (or nodes in the model) display correlated increases and decreases of power in specific carrier frequencies. Whole-brain network models - as the one presented herein - serve as ‘toy models’ to test different mechanistic scenarios where similar dynamical behaviours may be obtained with a reduced set of coupled dynamical equations. One way to validate the models is to quantify their fit with empirical data, for instance, by comparing the empirical and simulated frequency-specific FC matrices. Yet, comparing two models based on their quantitative fit of empirical data is not necessarily conclusive. For instance, even if the Hopf model reaches a fit of cc=0.5 and the Kuramoto reaches only a fit of cc=0.4 (Cabral et al., 2014), the main difference between the models does not rely on their *quantitative* performance in fitting the envelope FC, but rather on the fact that they are built on different assumptions.

In the model proposed in Cabral et al. (2014), local oscillations were restricted to the gamma-frequency range (i.e. with homogeneous frequencies at 40Hz). When coupled in the brain network, these gamma-band oscillators can temporarily synchronize with other brain areas at slower delay-dependent network frequencies falling in the alpha- and beta-frequency ranges (with a peak around 16Hz). This mechanism of metastable synchronization occurs in a critical working point where the system spontaneously switches between two regimes: one where the nodes oscillate at their fundamental frequency at 40Hz, and another where nodes synchronize with their neighbours at collective delay-dependent frequencies. However, this model builds on the assumption that brain areas are constantly engaged in self-sustained oscillations and their behaviour can be described by the Kuramoto model of coupled oscillators with time-delays. As such, it remains a theoretical scenario that needs further validation from experimental neurophysiological studies and/or from more detailed and biophysically realistic computational models.

In the model proposed herein, a brain area may be in a quiescent state and only engage in limit-cycle oscillations if the input is sufficiently strong to cross a supercritical bifurcation, as modelled by the normal form of a Hopf bifurcation. When embedded in a neuroanatomical network model, the input received from neighbouring areas modulates the amplitude of the oscillations. Since no mechanism of frequency reduction is implicit in this model, oscillations must be explicitly tuned in the alpha- and beta-frequency ranges, such that their power is modulated by the interactions in the neuroanatomical network. This builds on the assumption that sufficiently large brain areas may resonate at fundamental frequencies below the gamma-frequency range (note that gamma is so far considered the main resonant frequency of neural masses according to electrophysiological (Buhl et al., 1998) and theoretical studies (Brunel and Wang, 2003)). In addition, whether brain areas can resonate in parallel at a range of different frequencies – as assumed in the multi-frequency model – remains a purely conceptual scenario that requires further validation from the experimental/theoretical side. Yet, it serves to test a theoretical prediction before its empirical validation, forging new paths to investigate frequency-specific FC via computational models.

Importantly, rather than competing against each other for an optimal quantitative fit of MEG data, these modelling works are part of a collaborative effort to investigate the complex mechanisms underlying resting-state activity on multiple time-scales. Addressing the problem from different phenomenological perspectives, each model brings insightful information, irrespective of its individual flaws. In the complexity of the brain, it is not unlikely that the different mechanisms coexist and interplay at different levels, so these studies should be seen as complementary rather than exclusive, exploring new grounds that will eventually serve further experimental and computational studies. For instance, brain areas may not only resonate at multiple fundamental frequencies as proposed herein, but may additionally synchronize at myriad delay-dependent network frequencies if delays are considered.

### 4.3 Single-frequency Hopf Model

Following the line of previous whole-brain network models with locally-generated oscillations, we first considered a scenario where each brain area could generate oscillations in only one fundamental frequency. We tested a range of different fundamental frequencies and compared the results with the empirical data. We found that the best fit was obtained when the fundamental frequency matched the carrier frequency displaying the richest envelope dynamics in empirical data, namely 12 Hz. In this scenario, the amplitude of 12 Hz oscillations is modulated by the input received through the structural connectome. Areas belonging to a more densely coupled community receive correlated input and hence are more likely to display correlated amplitude envelopes. For this reason, we obtained a good fit of the envelope FC matrix for carrier bands around 12Hz. In addition, states of coherent envelopes (high synchrony) alternate with less coherent states, giving rise to strong fluctuations in the synchrony degree, similarly to what is observed in empirical MEG data. From the CCD analysis, we find similar time-dependencies because the patterns of envelope coherence (highly shaped by the structural connectivity) replicate over time. These results show that the complex envelope dynamics observed in resting-state MEG can be the signature of a relatively simple biophysical phenomenon where brain areas, composed of naturally excitable cortical tissue, are operating at the threshold of excitability. Although the model nicely fits the empirical observations in the most prominent frequency band, our results show that a single Hopf-model with an optimized fundamental frequency is not sufficient to explain the envelope dynamics in the remaining carrier frequency bands.

### 4.4 Multi-Frequency Hopf Model

To account for the limitations of the single-frequency model, we tested a conceptual model where each brain area could resonate simultaneously and independently at a range of fundamental frequencies. In more detail, each brain area consisted of several Hopf bifurcation models each with a different fundamental frequency. For each independent frequency layer, the nodes were coupled through the same structural connectivity (like in the single-frequency case). The selection of frequencies was purely phenomenological and, although it did not intend to match the peaks of the power spectrum typically found in resting-state recordings, it included values in the range of frequencies typically considered as resting-state carrier frequencies, namely the alpha and the beta bands (Hipp et al., 2012, Engel et al., 2013). This hypothetical scenario is supported by the fact that each brain area has complex internal and external connectivity patterns which may not unlikely resonate at a broad range of frequencies rather than a single one. Although the generative mechanisms of gamma-band oscillations have been subject of extensive study both in vivo, in vitro (Buhl et al., 1998) and through computational models (Brunel and Wang, 2003), one cannot neglect the clear evidence of oscillations in other frequency ranges (Berger and Gloor, 1969, Mantini et al., 2007, He et al., 2008, Scholvinck et al., 2010, Magri et al., 2012, Tagliazucchi et al., 2012, Keller et al., 2013), either arising (like gamma) from internal connectivity within a neural mass (Brunel and Wang, 2003), or from thalamo-cortical connectivity (Lopes da Silva et al., 1997), or even from delay and network-dependent frequencies defined by the large-scale neuroanatomical wiring structure (Cabral et al., 2014b).

As shown here, considering multiple rhythms within a brain area improved the performance of the model across the frequency spectrum. We had no intention to select an optimal number of frequencies, but rather to mimic a scenario where brain areas resonate at multiple fundamental frequencies in parallel, irrespective of their underlying generative mechanism. Recently, Hipp and colleagues (2015) found that a broad range of frequencies (2-128Hz) is implicated in resting-state functional connectivity, with different cortico-cortical connections being associated to distinct carrier frequencies. In the current implementation of the multi-frequency Hopf model, the same SC matrix shapes envelope correlations in all frequency layers. As such, little diversity of envelope FC matrices is obtained between frequencies, in contrast with what is observed in the empirical data (Hipp and Siegel, 2015). One way of overcoming this is to adjust the likelyhood that an area will resonate at each frequency by tuning the bifurcation parameter of each area *a_j_* for each fundamental frequency (e.g. informed by empirical MEG data). Importantly, what we learn from this model is that, unless delay-dependent network mechanisms generate more frequencies in the system (as in Cabral et al., 2014), multiple independent rhythm generators must be incorporated in future whole-brain network models in order to investigate the properties observed in resting-state MEG recordings at different frequency levels.

### 4.5 Cross-Frequency Interactions

In the multi-frequency model, we assumed *no* cross-frequency interactions. Actually, whether or not there is long-range coupling between different frequency bands in the brain remains under debate (Canolty and Knight, 2010, Engel et al., 2013, Aru et al., 2015). Indeed, even in the cases where experimental evidence of cross-frequency coupling is reported in the literature, either at the amplitude-to-amplitude (Furl et al., 2014, Brookes et al., 2016) or phase-to-amplitude level (Florin and Baillet, 2015), one cannot exclude the hypothesis that correlations between bands are generated by common biophysical processes unrelated to direct coupling or just by methodological pitfalls (Aru et al., 2015). Moreover, whilst there is extensive knowledge about the physiological mechanisms responsible for different frequency components (Buzsaki, 2006), not much is known about the cellular and network mechanisms of the *interactions* between these components (Canolty and Knight, 2010). Indeed, in order to validate the hypothesis of cross-frequency coupling in the brain, a biophysical theory needs to be put forward as to how a neuron or an ensemble of neurons physically implements the coupling. For example, Pastoll et al. (2013) have proposed a biophysical explanation to account for the theta-to-gamma (phase-to-amplitude) correlations, where the low frequency oscillation reflects periodic fluctuations of the membrane potential and thus excitability, which in turn gate the occurrence of rhythmic gamma oscillations. Yet, in the absence of proof to the contrary, assuming no cross-frequency interactions remains a plausible hypothetical scenario.

Considering no cross-frequency coupling at the generative level allows for an analogy with what is observed in networks of electromagnetic waves or radio waves. Indeed, frequency-division multiplexing is commonly used to transmit multiple independent signals over different non-overlapping carrier frequency bands (channels) via cables or optical fibres. In addition, this technique enables bidirectional communications over one strand of fibre, as well as multiplication of capacity (Weinstein and Ebert, 1971, Akam and Kullmann, 2014). As we show here, considering multiple uncoupled frequency generators at each brain area allows communication through independent frequency channels all sharing a common wiring structure of white matter pathways.

### 4.6 Amplitude vs Phase Coupling Modes

In this work, we focused in the amplitude coupling of MEG signals. This type of functional connectivity, also known as narrow-band envelope correlation or band-limited power modulation, operates on ultra-slow time-scales and has revealed meaningful resting-state networks similar to the ones observed in fMRI. Following the historical development of resting-state models, we focused on the genesis of slow amplitude modulations and neglected the faster interactions at the phase level. However, to understand long-range communication between brain areas requires deriving a complete picture of connectivity across coupling modes, including phase-to-phase, amplitude-to-amplitude and even phase-to-amplitude coupling (Engel et al., 2013). The model we propose here has the potential to be extended and optimized to serve as a tool to explore this multi-modal connectivity structure. Importantly, to focus on phase interactions rather than amplitude correlations, propagation times should be included in the model to account for non-negligible time-delays between distant brain areas.

## 5 Acknowledgements

GD is supported by the ERC Advanced Grant: DYSTRUCTURE (n. 295129) and by the Spanish Research Project PSI2013-42091-P. JC, AS, TVH and MLK were supported by the ERC Consolidator Grant CAREGIVING (n. 615539) and the TrygFonden Charitable Foundation. MWW is funded by the Wellcome Trust and an MRC UK MEG Partnership Grant (MR/K005464/1), and the National Institute for Health Research (NIHR) Oxford Biomedical Research Centre based at Oxford University Hospitals Trust.

